# Decoding central metabolic rewiring induced by exogenous GABA shunt intermediates in Pea

**DOI:** 10.64898/2026.07.28.741208

**Authors:** Ramnath Nayak, Portia D. Singh, Shyam Kumar Masakapalli

## Abstract

Graphical Abstract

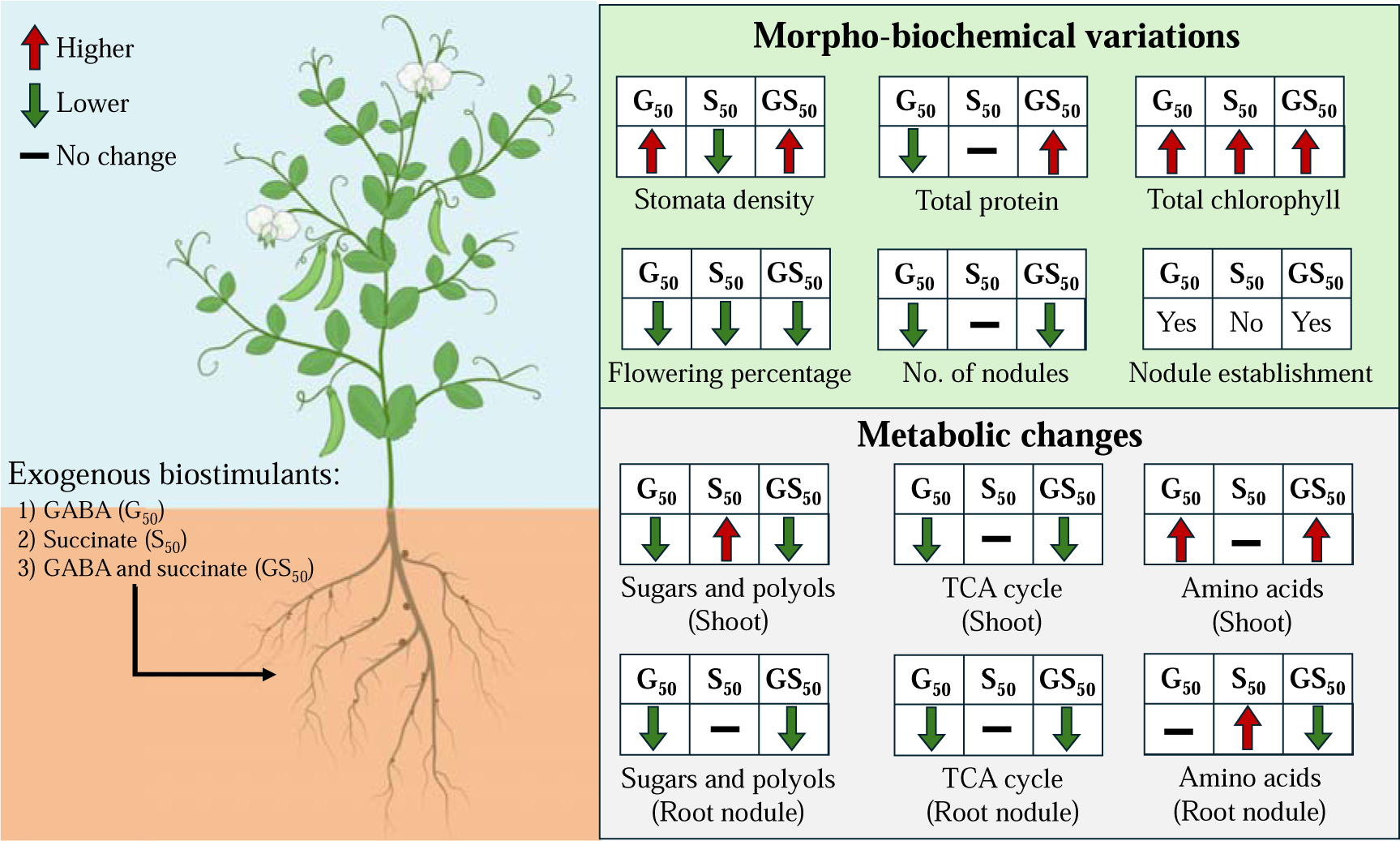

Legume-rhizobium symbiosis is constrained by carbon allocation and nutrient exchanges between the host plant and symbiotic bacteroids, emphasizing the necessity of biostimulant-based strategies to improve symbiotic efficiency. Studies suggest that plants provide carbon substrates (mainly TCA cycle intermediates) and nitrogen assimilation precursors to support bacteroid metabolism and nitrogen fixation. The GABA shunt represents a conserved bypass, linking the GS/GOGAT cycle, TCA cycle, and broader nitrogen metabolism. In the present study, the metabolic phenotypes upon rhizospheric application of exogenous γ-aminobutyric acid (GABA) and succinate were investigated in pea. Application of exogenous GABA and succinate primarily altered the morpho-biochemical properties and root nodule establishment of the plant. While GABA supplementation reduced the root length and increased stomatal density, succinate treatment alone inhibited nodule organogenesis with disrupted symbiosome zoning. Metabolic rewirings associated with these phenotypic changes begin at the root nodule compartment, where exogenous GABA altered endogenous tricarboxylic acid (TCA) cycle pools, leading to higher turnover of TCA cycle intermediates, including succinate, fumarate, and malate, as well as elevated levels of sugars (sucrose, glucose, and fructose) and polyols (pinitol, mannitol, and myo-inositol). Concurrently, the plants accumulated substantially higher levels of asparagine (+3.9 log_10_-fold higher), accompanied by altered shoot protein content and enhanced elemental nitrogen content (+0.8-1%). Collectively, these findings demonstrate the potential of GABA as a biostimulant capable of altering the carbon-nitrogen dynamics and enhancing symbiotic functioning in pea root nodules, whereas succinate alters the process of nodule formation.

## 1. Introduction

Legume root nodules are micro-anaerobic metabolic hubs where the host-symbiotic relationship thrives due to the associated metabolic exchanges and crosstalks (Chu et al., 2019; Lindström & Mousavi, 2020). The substantial energy demands of the symbiotic bacteroids are met by host plants mainly by channeling photosynthetic carbon derived from central metabolism, including glycolysis and the tricarboxylic acid (TCA) cycle, to the root nodule in exchange for fixed atmospheric dinitrogen (Desbrosses & Stougaard, 2011; Vacheron et al., 2013). However, only a minimal fraction of photosynthates reaches the root nodule, representing a major constraint to nodule activity (Hennion et al., 2019; Shabala et al., 2016). To address this limitation, there is growing interest in identifying and utilizing potential biostimulants to mitigate the constraints on plant-microbe symbiosis. Exogenous application of various biostimulants has been reported to improve nutrient-use efficiency (Canellas et al., 2015; Halpern et al., 2015; Li et al., 2024; Paul et al., 2019), enhance tolerance to abiotic stresses (da Silva et al., 2023; El-Nakhel et al., 2023; Lucini et al., 2015), and regulate major metabolic processes (Chakrabarti & Mukherji, 2003). Unlike supernodulating genotypes, which often incur additional carbon costs for nodule maintenance and may compromise biomass production (Zhong et al., 2024), biostimulants have the potential to improve symbiotic performance by providing supplementary metabolic support to the plant and driving significant metabolic reprogramming (Batushansky et al., 2014). Despite these promising attributes, the metabolic mechanisms underlying biostimulant-driven responses in legume symbiosis remain insufficiently characterized.

Among the metabolites involved in legumes’ C/N homeostasis, γ-aminobutyric acid (GABA) and succinate play key roles, connecting carbon metabolism (TCA cycle) and nitrogen metabolism (glutamine synthetase-glutamate synthase [GS–GOGAT] cycle) through the GABA shunt pathway. This is a three-step bypass that uses glutamate decarboxylase (GAD), GABA transaminase (GABA-T), and succinic semialdehyde dehydrogenase (SSADH) to convert glutamate into succinate, providing an alternative route for carbon to enter the TCA cycle (Fait et al., 2008). GABA is a non-proteinogenic amino acid that accumulates during nodule development and functioning (Sulieman & Schulze, 2010), contributing to carbon-nitrogen balance (Jurgen et al., 2009) and stress adaptation (Nikhil et al., 2023; Xu et al., 2021; Yong et al., 2017). Succinate, a central intermediate of the TCA cycle, serves as an important carbon and energy source for bacteroid metabolism inside root nodules (Schulte et al., 2025; Terpolilli et al., 2016). Beyond its metabolic role, GABA functions as a signaling molecule with key roles in nutrient utilization (Wang et al., 2025), cellular signaling (Bown & Shelp, 2016; Suhel et al., 2023; Xu et al., 2021), and metabolic plasticity (Chen et al., 2022; Yong et al., 2017). Given the tightly coordinated carbon and nitrogen metabolism in legumes, both GABA and succinate have emerged as promising metabolic biostimulants capable of modulating metabolic homeostasis and adaptive physiological responses (Sulieman et al., 2024).

Although GABA and succinate are recognized as central intermediates necessary for host-symbiont survival, their role as potential metabolic biostimulants in reorganizing the global metabolome remains poorly understood. The majority of existing studies have focused on their individual physiological functions, while comparatively little attention has been directed to how exogenous supplementation reshapes global metabolic networks. Furthermore, whether GABA and succinate elicit distinct, overlapping, or synergistic metabolic responses when applied in combination remains uncertain (Sulieman, 2011). Addressing these gaps is essential for a more profound understanding of how GABA shunt intermediates sustain and support symbiotic root nodule function under varying metabolic demands.

Pea (*Pisum sativum* L.) is an agriculturally important legume with active carbon and nitrogen metabolism and a strong dependence on coordinated metabolic regulation to support growth and symbiotic function. Unlike model legumes such as *Medicago truncatula* or *Lotus japonicus*, pea offers the agronomic advantage of being a major food and feed crop in temperate regions, making insights into its symbiotic metabolism directly translatable to sustainable agriculture. We hypothesized that applying GABA and succinate in the rhizosphere, both separately and together, would cause distinct but overlapping changes in metabolism in the shoot, nodulated roots, and rhizosphere compartments, improving nitrogen assimilation by altering the carbon-nitrogen crosstalk. In this study, we characterized changes in metabolic phenotypes upon rhizospheric application of GABA shunt intermediates (GABA and succinate) in pea-microbe interactions by: (1) examining the effects on plant growth (stomatal density and flowering), root architecture, and nodule structure; (2) assessing changes in the nitrogen status of peas (protein content and CHNS elemental analysis); and (3) performing comprehensive metabolic profiling of shoot, nodulated root, and rhizosphere metabolites using GC-MS and ¹H-NMR. By characterizing the tissue-specific changes in amino acids, organic acids, sugars, and associated metabolic networks, this study aims to decipher the metabolic rewiring associated with biostimulants and offer novel insights into their roles in regulating plant metabolism in support of pea-microbe symbiosis.

## 2. Materials and Methods

### 2.1 Plant material, rhizobium inoculation, and growth conditions

Pea seeds (*Pisum sativum* var. AS-10) were surface-sterilised using 0.1% sodium hypochlorite solution for 5 min, followed by washing with sterile water. After washing, seeds were germinated in the dark at 25°C for 4 days in a soil mixture of coco peat and vermiculite (1:1, w/w). No rhizobium inoculation was performed, and the nodulation was due to natural infection by indigenous rhizobial populations present in the field soil at IIT Mandi, Himachal Pradesh, India. Soil rhizobial community composition was not characterized in the present study. Post-germination, seedlings (3 per pot) were transplanted into earthen pots (20×10×10cm). Eight replicates were used per treatment. Pots were relocated to field conditions at IIT Mandi (31°46′28″ N, 76°59′10″ E), India, in August 2024. Temperature fluctuations ranged from a maximum of 36.1°C to a minimum of 16.6°C (average 24.8°C), with relative humidity ranging from 40.8% to 99.2%, recorded using a Tempnote (TH32) device. Pots were regularly rotated to ensure uniform light exposure.

### 2.2 Application of biostimulants

50 mM concentration of exogenous GABA, succinate, or a combination of both was formulated for rhizospheric application. The following abbreviations are used throughout:

- **C_0_ 0DAT:** No treatment control, harvested on the day of treatment
- **C_0_ 10DAT:** No treatment control, harvested 10 days after treatment (DAT)
- **G_50_ 10DAT:** 50 mM GABA treatment, harvested 10 DAT
- **S_50_ 10DAT:** 50 mM succinate treatment, harvested 10 DAT
- **GS_50_ 10DAT:** 50 mM GABA + 50 mM succinate, harvested 10 DAT

### 2.3 Sample harvesting and morphological characterization

The lengths of shoots and roots were calculated using ImageJ software (https://imagej.net/ij/). Relative ratios were considered to minimise biological variation. The moisture content of the samples was calculated using the formula:

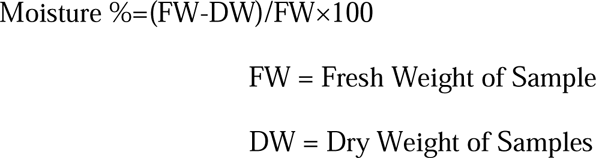

where FW = fresh weight and DW = dry weight. Plant shoots and nodulated roots (n = 4 biological replicates) were separated using a razor blade and harvested at two time points (days 20 and 30) by quenching in liquid N_2_ followed by lyophilisation. Samples were finely powdered using a mortar and pestle and stored at −20°C for biochemical and metabolomic analysis. Soil samples were collected from within a 5 cm radius of pea roots at days 0, 20, and 30 and stored for biochemical analysis.

### 2.4 Tissue staining and microscopic analysis

Stomatal distribution along the first leaf was examined using the nail polish imprint technique (Pathoumthong et al., 2023). Ten days post-treatment, leaf imprints were obtained by staining the abaxial surface with 0.1% safranin and mounting on glass slides. Images were acquired using a Nanosight microscope (MALVERN) at 13.5× magnification, and stomatal density (mm^-2^) was calculated. One-way ANOVA was used to assess significant differences.

For visualization of indeterminate pea root nodules, freshly harvested roots were washed in PBS containing 50 mM PIPES buffer (pH 7.0). Nodules were categorized as normal (N) or altered (AN) based on morphology. Nodules were fixed in 4% paraformaldehyde for 10 min, sectioned with a double-edged razor blade, and stained with 1 μL/mL Hoechst (Thermo Fisher Scientific, R37605) for 15 min. Confocal images were acquired in the DAPI channel using a Nikon ECLIPSE Ti2 inverted microscope system (Nikon Instruments Inc., Melville, NY). Fluorescence was displayed in green; images were processed using ImageJ.

### 2.5 Determination of total proteins, chlorophyll, and carotenoid content

Total protein was extracted and quantified from 10 mg lyophilized material as described in (Pant et al., 2023). Briefly, proteins were extracted in 200 μL extraction buffer, vortexed for 20 s, and incubated on ice for 5 min (repeated three times for 15 min total), followed by centrifugation at 12,600 × g for 20 min at 4°C. The supernatant was collected, and protein content was quantified using the DC Protein Assay (Bio-Rad, catalog no. 500-0116) at 750 nm against a BSA standard curve. Elemental composition (C, H, N, S) was analyzed using the UNICUBE organic elemental analyzer (Elementar, Germany). For chlorophyll and carotenoid determination, 100 mg fresh leaves were crushed in DMSO and centrifuged at 10,000 × g; 200 μL of supernatant was measured at 647 and 664 nm, and pigment content was calculated as described in Pant et al. (2023).

### 2.6 Metabolite extraction and metabolomic analysis

Polar metabolites were extracted from 20 mg freeze-dried plant material in 80% (v/v) aqueous methanol by boiling at 70°C at 950 rpm for 5 min (repeated twice), followed by centrifugation at 13,000 × g for 10 min at room temperature. Aliquots were vacuum-dried for NMR analysis. NMR extracts were dissolved in 600 μL of deuterated water that contained 1 mM trimethylsilyl-2,2,3,3-tetradeuteropropionic acid sodium salt (TSP, 0.01% v/v), 1 mM EDTA, and 50 mM KH_2_PO_4_. Spectra were acquired with 128 scans on a 500 MHz JEOL ECX500 spectrometer (acquisition time 1.75 s; relaxation delay 4 s; pulse angle 45°; pulse width 1.75 μs) and analyzed using JOEL Delta, MestReNova, and Chenomx NMR Suite.

For GC-MS, metabolites from 20 mg plant material were extracted in 1,740 μL cold chloroform:methanol:water (3:1:1 v/v/v) (Lisec et al., 2006). Rhizosphere metabolites were extracted from 1 g soil using 1 mL aqueous methanol (1:1, v/v). Ribitol (0.2 mg/mL) was used as the internal standard. Fifty μL of each extract was dried in a SpeedVac, then derivatized with methoxyamine hydrochloride (MeOX) and trimethylsilyl (TMS) reagent (Singh et al., 2026). Derivatized samples (1 μL) were analyzed in splitless mode on a GC-MS system (GC ALS-MS 5977B, Agilent Technologies) equipped with an HP-5ms column (30 m × 250 μm × 0.25 μm) at IIT Mandi. Helium carrier gas was used at 0.6 mL/min over a 60-min run. The oven program started at 50°C, ramped to 70°C with a 5-min hold, then increased at 10°C/min to 200°C with a 10-min hold, and finally ramped at 5°C/min to 300°C with a 10-min hold. Detection was performed over m/z 50-600 at 70 eV electron ionization. Peak identification used the NIST 17 library (probability > 75%), with spectral alignment via MetAlign. Statistical and multivariate analyses were performed in MetaboAnalyst v6.0 (https://www.metaboanalyst.ca/). Data were normalized to the internal standard ribitol to calculate relative metabolite abundances and fold changes. All metabolite abundance values are reported as relative abundances normalized to the internal standard and should be interpreted accordingly.

## 3. Results

### 3.1 Exogenous GABA and succinate have differential effects on pea growth, physiology and nodule morphology

To capture the phenotypic changes upon the rhizospheric application of the treatments (50 mM GABA and/or succinate), morpho-biochemical responses of the pea plants were evaluated across two developmental stages: nodule organogenesis and flowering, following treatment at day 20 (0 DAT) (Figure S1). The key changes in the metabolic phenotype of the plants were captured during the 0-10 DAT window (days 20 and 30), by agronomic measurements, biochemical assays, and metabolite profiling of the shoot and nodulated root tissues.

Treatment with 50 mM GABA (G_50_) significantly altered shoot-to-root biomass allocation, resulting in reduced root mass and root length, relative to control plants (Figure 1A and 1B, Table 1). Stomatal density was significantly increased in GABA-treated plants (G_50_ and GS_50_) (Figure 1C). Assessment of nitrogen status revealed divergent protein responses between treatments: G_50_ significantly reduced shoot protein content (−Δ 16.5 mg g^-1^ FW), whereas GS_50_ elevated it (+Δ 20 mg g^-1^ FW) (Figure 1D). Elemental nitrogen (N%) was significantly increased exclusively under G_50_ treatment (Table 1). Total chlorophyll and carotenoid contents were significantly elevated across all treatments relative to controls (Figures 1F-1G).

**Figure 1:**
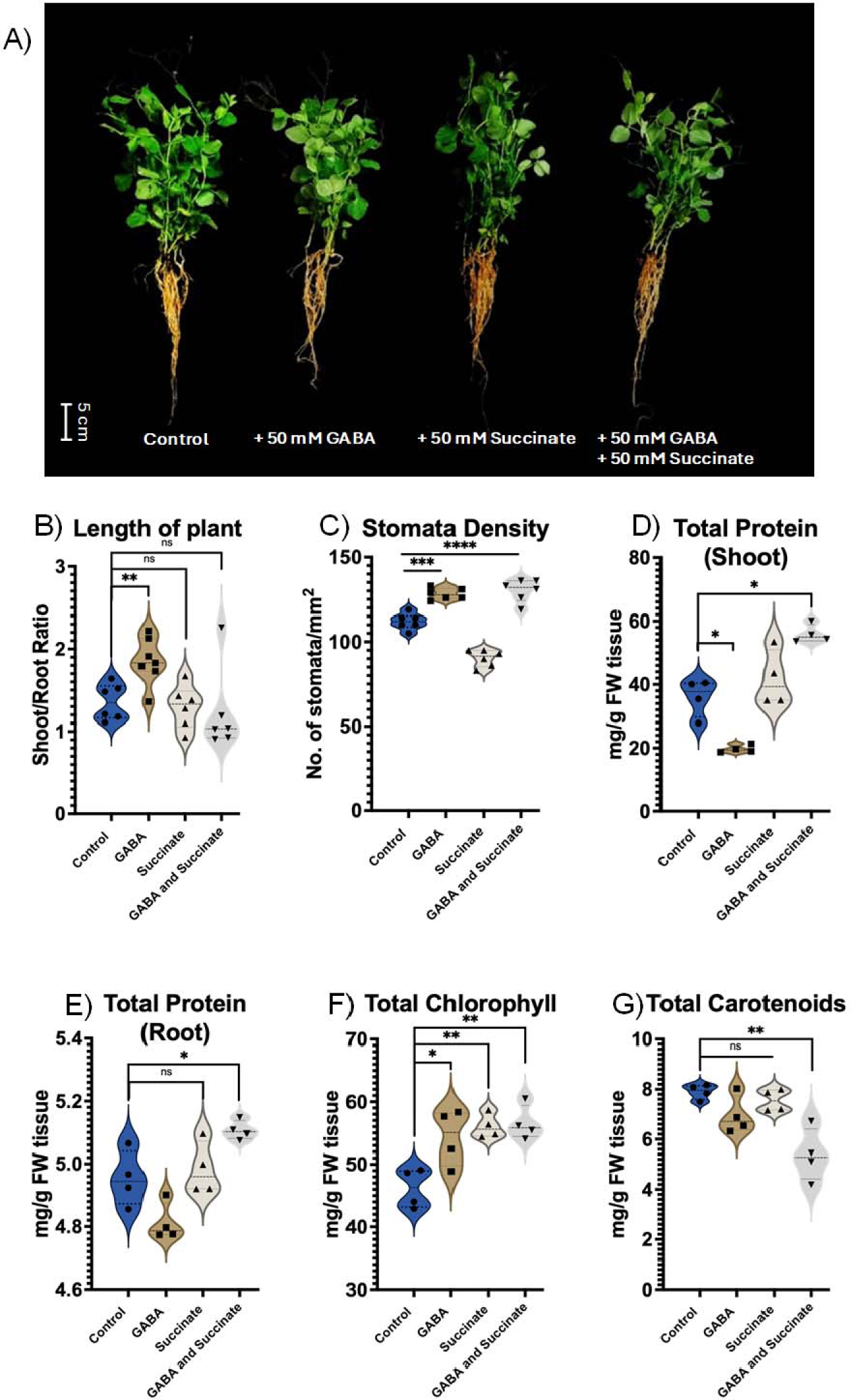
Morpho-biochemical properties of pea plants subjected to exogenous GABA and/or succinate. A) 30-day-old pea plants subjected to exogenous a) control (C0), b) 50 mM GABA (G50), c) 50 mM succinate (S50), and d) 50 mM GABA and 50 mM succinate (GS50). B-C) shows the variations in the ratio of shoot: primary root length and the changes in the stomata density, respectively. D-E) shows the differences in the total protein content in the shoot and root, respectively. F-G) shows the changes in the photosynthetic pigments, chlorophyll, and carotenoid. All plants were treated with/without exogenous GABA and/or succinate. The data is presented using a violin plot showing the distribution of various parameters. One-way ANOVA followed by Tukey’s HSD post-hoc test was used to find the significant differences between the groups. Asterisks denote the statistical variations between treatments. The data is presented using a violin plot (*p < 0.5, **p < 0.05, and ***p < 0.005).

**Table 1.**
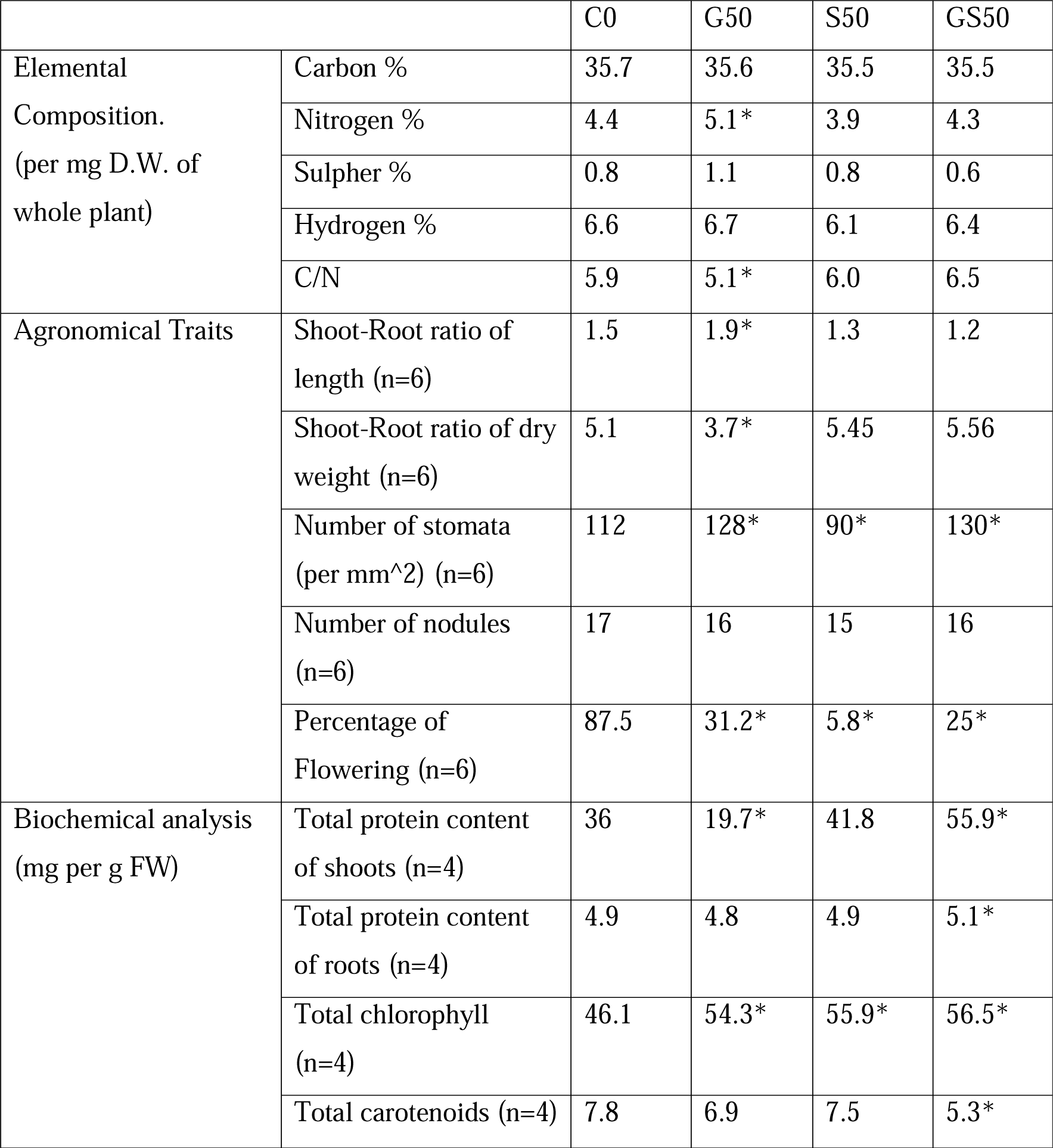
The morphological variations observed 10 days after treatment in the pea inoculated with the biostimulants. * represents statistically significant values (p<0.5) analysed using One-way ANOVA. The abbreviations are: control (C0), 50 mM GABA (G50), 50 mM succinate (S50) and 50 mM GABA+succinate (GS50).

Rhizospheric application of succinate (S_50_ and GS_50_) induced substantial alterations in nodule morphology and structure across both primary and lateral root systems. Based on the phenotype, the nodules were categorized as normal (N) or altered (AN) (Figure 2A). Altered nodules exhibited marked loss of characteristic pink pigmentation relative to normal nodules (Figure 2B). Histological analysis of altered nodules revealed disrupted nodule zonation, characterized by impaired symbiosome differentiation within zone III (Figure 2C). Despite these structural alterations, moisture content of nodulated roots was significantly increased under G_50_ and S_50_ treatments (Figure 2D). No significant differences were observed in nodulated root dry mass; however, nodule number was significantly reduced in GABA-treated plants (G_50_ and GS_50_) (Figures 2E-2F).

**Figure 2:**
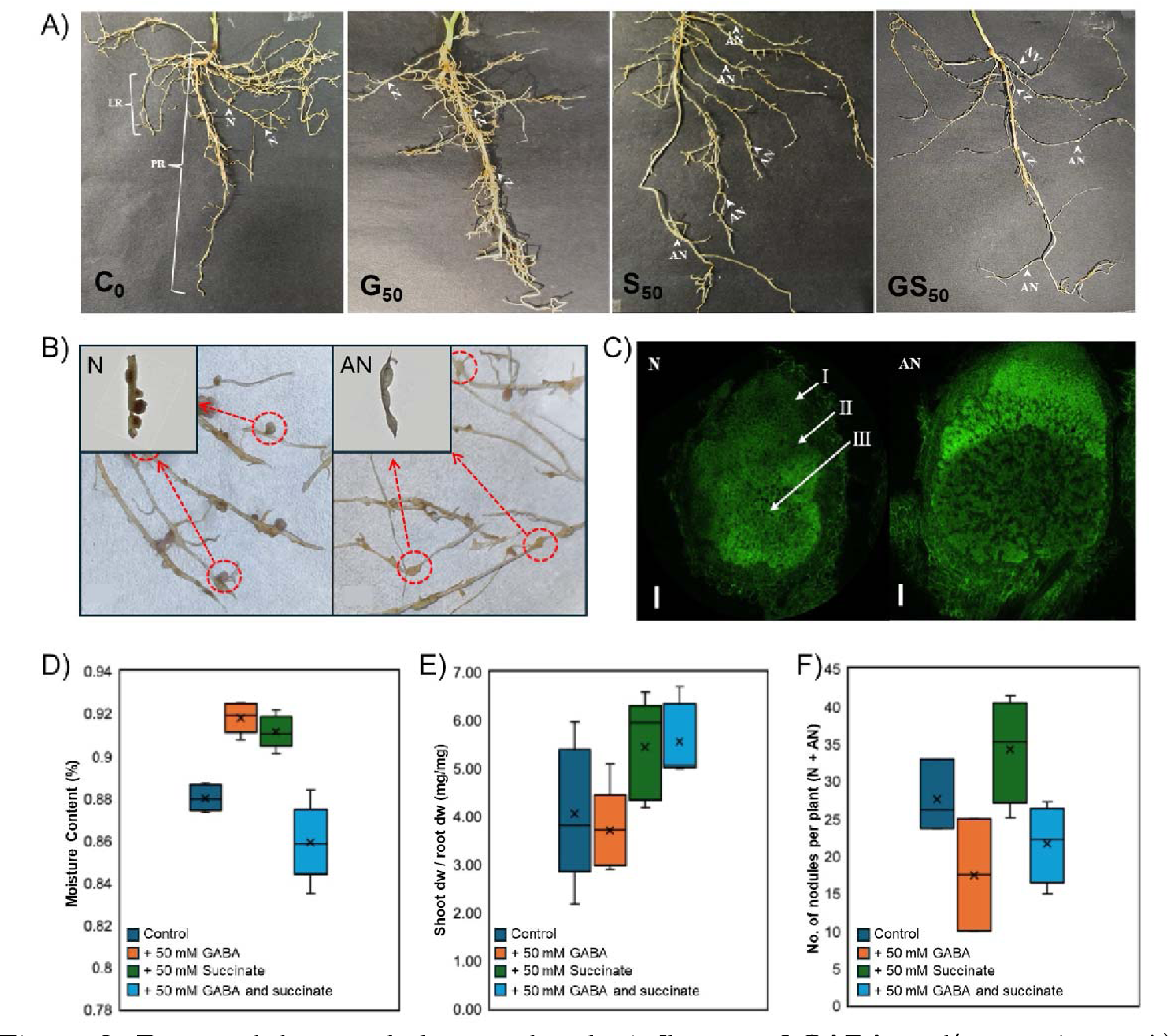
Root-nodule morphology under the influence of GABA and/or succinate. A) Shows the changes in the root and nodule morphology that were observed 10 DAT in the primary (PR) and lateral (LR) root structure upon the administration of exogenous treatments of 50 mM GABA (G_50_), 50 mM succinate (S), and 50 mM GABA and 50 mM succinate (GS_50_). B) Alterations in nodule morphology. The intact structure (pink) of the nodule is denoted by ‘N’ whereas the altered nodule is denoted by ‘AN’. C) Confocal images of the nodule structures N and AN. The nodule zones (Zone I-III) are depicted here. Bar scale: 10 μm. D-F) Box plot representing the morphological status of roots by considering moisture content, dry weight ratio of shoot and root, and number of nodule structures (N + AN) per plant, respectively.

### 3.2 Untargeted metabolomics highlights distinct tissue-specific metabolic adjustments induced by rhizospheric GABA and succinate application

To characterize tissue-specific metabolic changes, untargeted metabolomics was performed using GC-MS (Figures 3A and 4A) and ^1^H-NMR (Supplementary Figure S3) on shoot and nodulated root tissues at 0 and 10 DAT. GC-MS profiling identified 60 metabolites in shoot tissue and 45 metabolites in nodulated root tissue (Supplementary Tables S2 and S3). These metabolites were categorized into five functional classes: (1) TCA cycle intermediates, (2) sugars and polyols, (3) nitrogen-based metabolites, (4) organic acids, and (5) other miscellaneous compounds.

**Figure 3:**
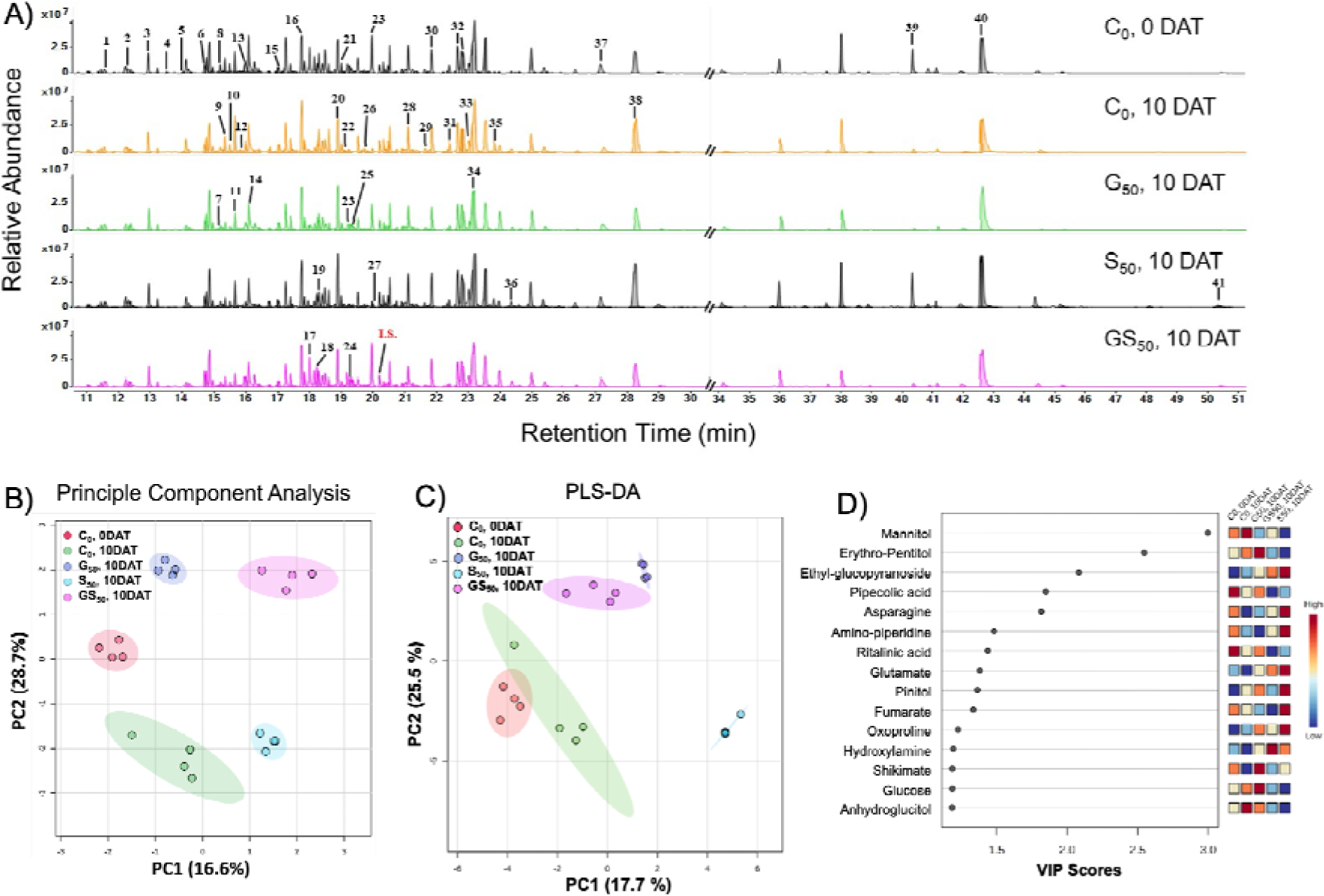
Metabolic variations observed in the nodulated pea shoot treated with GABA and/or succinate. A) GC-MS spectra of the soluble extracts of pea shoots capturing the 41 metabolites (see Supplementary table S2). IS is the internal standard ribitol (0.01% w/v). B-C) Metabolic variations in the shoot upon exogenous treatment, captured by principal component analysis and PLS-DA of the samples. The shaded regions highlight 95% confidence intervals between the treatments. D) Variable importance in projection (VIP) scores highlight the top 15 important metabolites in the shoot. The colors in the VIP scores represent high (red) to low (blue) values. The analysis was conducted in biological replication (n=4). C0, 0 DAT – control plants without any treatment at day 0; C0, 10DAT – control plants without any treatment at day 10; G50, 10DAT –plants treated with 50 mM GABA at day 10; S50, 10DAT –plants treated with 50 mM succinate at day 10; GS50, 10DAT –plants treated with 50 mM GABA and 50 mM succinate at day 10.

**Figure 4:**
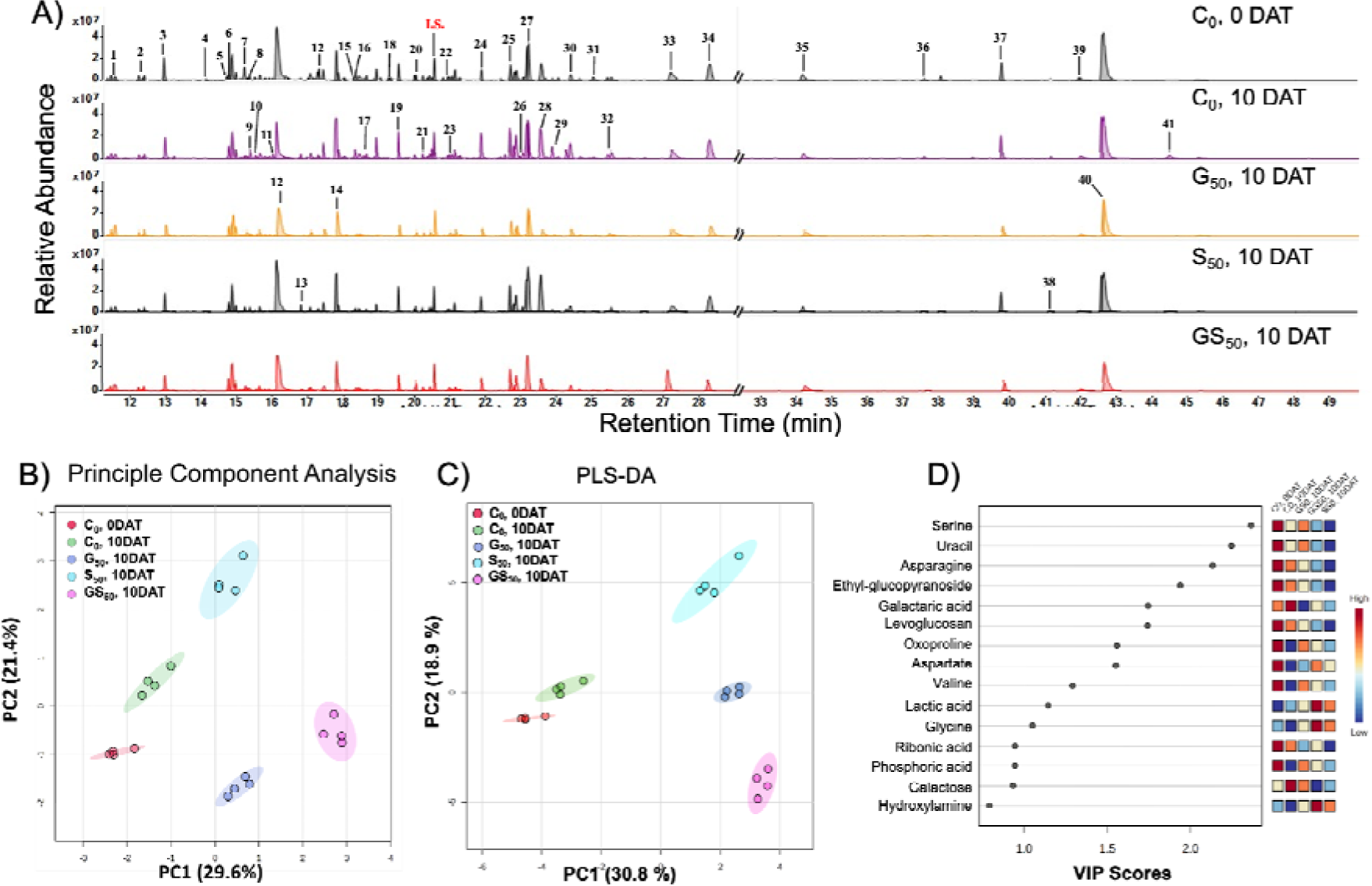
Metabolic variations observed in the nodulated pea roots (NPR) treated with GABA and/or succinate. A) GC-MS spectra of the soluble extracts of NPR capturing the 41 metabolites (see Supplementary table S3). IS is the internal standard ribitol (0.01% w/v). B-C) Metabolic variations in the NPR upon exogenous treatment, captured by principal component analysis and PLS-DA. The shaded regions highlight 95% confidence intervals between the treatments. D) Key metabolites contributing to the variations in NPR subjected to exogenous treatments are derived from variable importance in projection (VIP) scores. The colors in the VIP scores represent high (red) to low (blue) values. The analysis was conducted in biological replication (n=4). C0, 0 DAT – control plants without any treatment at day 0; C0, 10DAT – control plants without any treatment at day 10; G50, 10DAT –plants treated with 50 mM GABA at day 10; S50, 10DAT –plants treated with 50 mM succinate at day 10; GS50, 10DAT –plants treated with 50 mM GABA and 50 mM succinate at day 10.

Principal component analysis (PCA) revealed clear metabolic separation between treatments. In shoot tissue, PC1 and PC2 accounted for 16.6% and 28.7% of total variance, respectively (combined: 45.3%), whereas in nodulated root tissue, PC1 and PC2 explained 29.6% and 21.4%, respectively (combined: 51%) (Figures 3B and 4B). G50 and GS50 treatments exhibited distinct metabolic profiles relative to controls and S50 along both principal components. PLS-DA with Variable Importance in Projection (VIP score > 1.5) identified the metabolites most contributing to group separation (Figure 3C-3D, Figure 4C-4D). In shoot tissue, the primary discriminating metabolites were mannitol, erythron-pentitol, ethyl-glucopyranoside, pipecolic acid, and asparagine (Figure 3D). In nodulated root tissue, serine, uracil, asparagine, ethyl-glucopyranoside, and galactaric acid were the principal contributors to group discrimination (Figure 4D).

Rhizosphere metabolite profiling by GC-MS identified 35 metabolites, representing the small-molecule pool present at the root-soil interface during symbiosis (Figure S4). Multivariate analysis of rhizosphere profiles revealed treatment-specific clustering, primarily driven by differential levels of hydroxybutyric acid, homoserine, sucrose, and maltose. Cross-platform comparison of GC-MS and ^1^H-NMR data demonstrated consistent patterns for key metabolites, including asparagine and malate (Supplementary Figure S3), confirming the major metabolic changes identified across both analytical platforms.

### 3.3 Exogenous GABA reshapes the carbon-nitrogen metabolism in the pea nodule system

To contextualize the metabolic response, metabolomic profiling was first performed under non-symbiotic (NS) conditions (Supplementary Figure S2). Under NS conditions, PCA revealed distinct separation between control and GABA-treated samples in both tissues (Supplementary Figures S2A, S2C). Shoot metabolic divergence was driven by proline and tryptophan (VIP > 2), while fumarate and galactinol were the primary discriminating metabolites in roots (Supplementary Figures S2B, S2D). Heatmap analysis revealed a marked decline in nitrogenous metabolites in shoot tissue under NS conditions. In contrast, root tissue accumulated TCA cycle intermediates, including fumarate, succinate, citrate, and malate, under NS conditions.

Under symbiotic conditions, exogenous GABA induced distinct metabolic reprogramming characterized by coordinated redistribution of carbon and nitrogen metabolites between shoot and nodulated pea roots (NPR) (Figure 6). In NPR, increased turnover of carbon reserves was observed, including sugars such as sucrose, glucose, fructose, mannose, myo-inositol, and pinitol, alongside depletion of TCA cycle intermediates: citrate (−0.64 log_10_-fold relative abundance), succinate (−0.18 log_10_-fold relative abundance), fumarate (−2.59 log_10_-fold), and malate (−0.32 log_10_-fold relative abundance).

Depletion of these carbon pools was accompanied by substantial accumulation of nitrogen-rich metabolites in shoot tissue, most notably asparagine (+3.9 log_10_-fold relative abundance) (Figure 6). GC-MS and ^1^H-NMR data showed consistent patterns for asparagine across both platforms (Supplementary Figure S3). Shoot metabolic changes also included elevated glutamate (+0.56 log_10_-fold relative abundance) and decreased pipecolate (−0.7 log_10_-fold relative abundance). Collectively, these data demonstrate that rhizospheric application of 50 mM GABA induced coordinated depletion of root carbon reserves and accumulation of nitrogen-rich metabolites in shoot tissue under symbiotic conditions.

## 4. Discussions

The energetic demands of symbiotic bacteroids impose a substantial carbon cost, which the host plant meets by redirecting approximately 3% of photosynthates to root nodules (KOUCHI et al., 1986; Libault, 2014; Minchin & Witty, 2005) in exchange for fixed nitrogen supplied by bacteroids. Within the nodule, the host provides TCA dicarboxylates, primarily malate and succinate, in exchange for symbiotically fixed ammonia, which is subsequently assimilated via the GS/GOGAT pathway using host-supplied carbon substrates (Liu et al., 2018; Pfau et al., 2018; Ronson et al., 1981; Udvardi & Poole, 2013). A portion of the fixed nitrogen is utilized by root cells, while the remainder is transported to the shoot primarily as asparagine, owing to its favorable C/N ratio and stability during long-distance phloem transport (Parsons & Sunley, 2001; Rosa-Téllez et al., 2026; Sulieman & Tran, 2013). This study provides insight into how exogenous GABA shunt intermediates alter metabolic phenotypes in pea-microbe interactions.

A key observation in the present dataset is the contrasting metabolic response to GABA under non-symbiotic versus symbiotic conditions. Under non-symbiotic conditions, exogenous GABA treatment led predominantly to accumulation of TCA cycle intermediates in roots without a corresponding enhancement of nitrogen assimilation in shoot tissue, indicating that GABA functions primarily as a source of carbon skeletons rather than as a nitrogen source under these conditions (Supplementary Figure S1). Under symbiotic conditions, however, GABA treatment was associated with greater carbon depletion in root nodules alongside pronounced accumulation of the asparagine pool in shoot tissue (Figures 5 and 6). These observations suggest that exogenous GABA acts through the mitochondrial GABA shunt, funneling carbon into the TCA cycle via succinate formation, which in turn sustains bacteroid respiration and supports nitrogen fixation energetics via the malate-to-pyruvate shunt, mediated by NADL-dependent malic enzyme (Schulte et al., 2025; Terpolilli et al., 2016).

**Figure 5:**
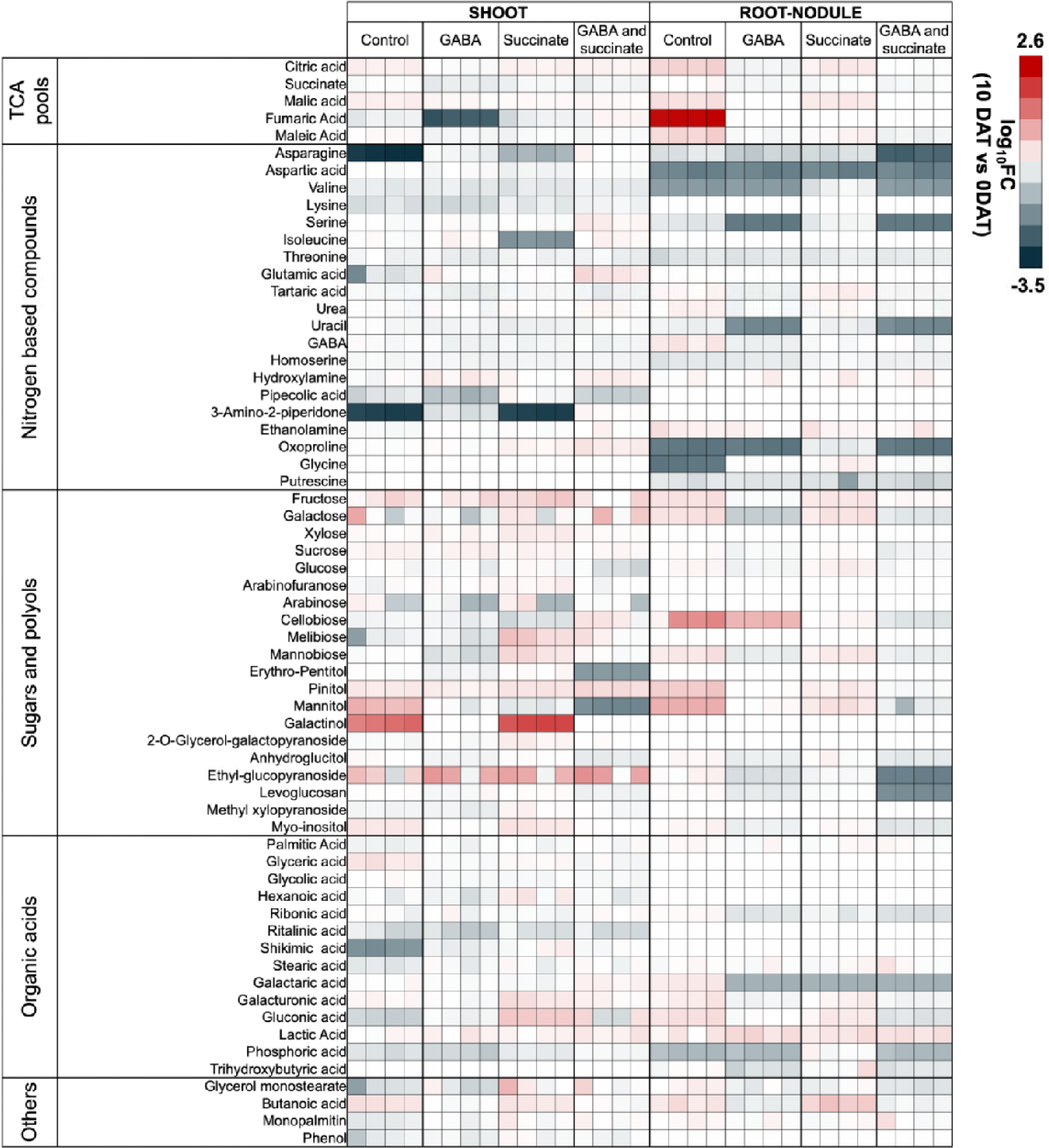
Metabolic changes in the pea shoot and root nodule post-biostimulant treatment are signified by the heatmap. The altered global metabolism of shoot and root nodules in pea treated with exogenous biostimulants (G_50_, S_50_ and GS_50_) (n=4). The values highlight the log_10_ fold-change values (10 DAT vs. 0 DAT) of different classes of compounds that were captured. The color of each metabolite indicates the class of compounds they belong to.

**Figure 6:**
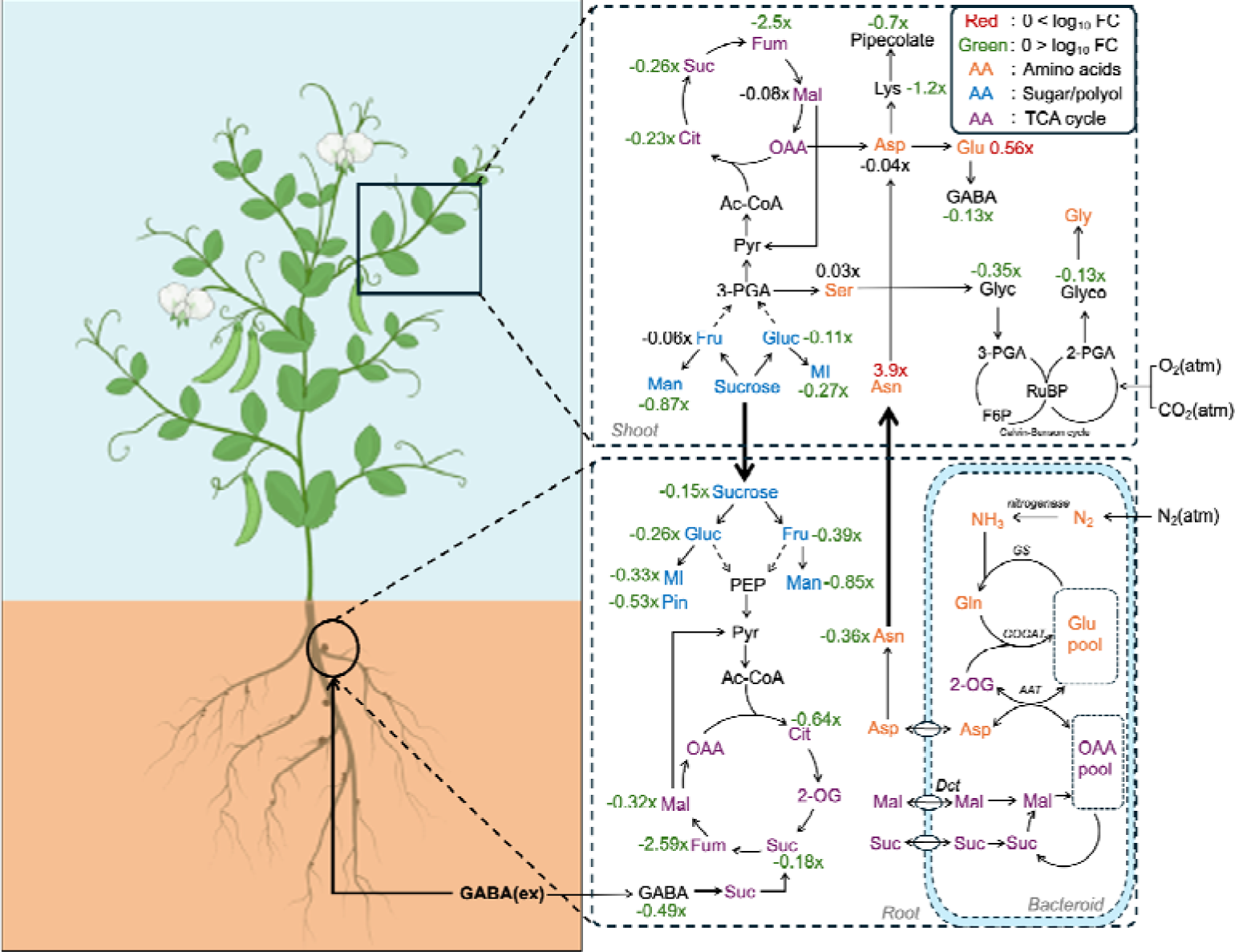
Exogenous GABA induces central metabolic rewiring and altered carbon-nitrogen partitioning in pea. Rhizospheric application of 50 mM exogenous GABA modulates the root and nodule metabolic crosstalk by altering the levels of sugar allocation, amino acid metabolism, and tricarboxylic acid (TCA) cycle intermediates. The reduced levels of sugars/polyols and several TCA intermediates suggest increased metabolic turnover and respiratory utilization to sustain symbiotic activity. This in return promotes higher nitrogen capture, as reflected by the elevated asparagine levels in the shoot, which indicates enhanced nitrogen export from root nodules to aerial tissues. Metabolites shown in orange represent amino acids, blue represents sugars/polyols, and purple represents TCA-cycle intermediates. Red values indicate increased metabolite abundance, whereas green values indicate decreased abundance relative to control plants. The depicted values are in terms of log 10-fold changes w.r.t. control. Green and red values indicate metabolites whose levels were either decreased or elevated, respectively. Metabolites whose levels were unchanged are marked by black values. Gluc: glucose, MI: myo-inositol, Pin: pinitol, Fru: fructose, Man: mannitol, PEP: phosphoenolpyruvate, 3-PGA: phosphoglyceric acid, Pyr: pyruvate, Ac-CoA: acetyl-CoA, Cit: citrate, 2-OG: 2-oxoglutarate, Suc: succinate, Fum: fumarate, Mal: malate, OAA: oxaloacetate, GABA: gamma-aminobutyric acid, Asp: aspartate, Asn: asparagine, Glu: glutamate, Gln: glutamine, Lys: lysine, Glyc: glycerate, Glyco: glycolate, GS: glutamine synthetase, GOGAT: glutamate synthase, AAT: aspartate aminotransferase, and Dct: dicarboxylate transporter.

The substantial increase in shoot asparagine levels observed under exogenous GABA treatment (+3.9 log_10_-fold relative abundance) represents a central finding of this study. This accumulation may be attributable to three non-mutually exclusive mechanisms: (1) enhanced de novo asparagine synthesis via asparagine synthetase (AS), driven by increased glutamine and oxaloacetate availability from an active GS/GOGAT-TCA cycle, a well-characterized route in leaves (Canales et al., 2012; Qiao et al., 2026); (2) reduced asparagine catabolism (Yabuki et al., 2017); and (3) increased phloem loading and export from root nodules (Sulieman et al., 2010; Sulieman & Schulze, 2010; Sulieman & Tran, 2013). The elevated elemental N% in GABA-treated plants (Table 1), together with ^1^H-NMR-confirmed asparagine accumulation (Supplementary Figure S3), supports the interpretation that this response reflects genuine nitrogen assimilation. However, the concentration of GABA applied (50 mM), which substantially exceeds endogenous tissue levels, warrants caution in extrapolating these findings to field conditions; future dose–response studies are therefore necessary. The apparent discrepancy between reduced shoot protein content under G_50_ and elevated N% may reflect preferential partitioning of fixed nitrogen into free amino acid pools, particularly asparagine, rather than into structural protein synthesis (Dokwal et al., 2022). Although the metabolic profiles of G_50_ and GS_50_ treatments showed overall similarity (Figures 3B and 4B), contrasting effects on protein content and N% were observed between treatments. Under combined GABA and succinate application, shoot protein content was elevated without a corresponding increase in N%, suggesting differential regulation of nitrogen partitioning between the two treatments.

Given the central role of succinate in nodule metabolism, one might expect that exogenous priming would promote the nodule activity. However, the altered nodule phenotype (Figure 2) showed an opposite trend, offering a newer insight into how exogenous succinate priming alters nodule zoning. Metabolite profiling suggests that, in contrast to GABA treatments, exogenous succinate (S_50_) primarily altered the metabolite profiles in the root system, having elevated levels of sugars (sucrose, glucose and fructose) and polyols (pinitol, mannitol and myo-inositol), while soluble nitrogen pools remained largely unperturbed, except for the levels of serine and ethanolamine (Figure 5), indicating that a failed nodule establishment caused a significant redistribution along glycolytic arms (Figure 7). Additionally, the elevated rhizosphere levels of sugars (arabinose, fructose, glucose, galactose, lactose, maltose, and sucrose) and sugar alcohols (mannitol and myo-inositol) observed under succinate treatment (Supplementary Figure S4) suggest a stimulation of soil microbial activity by succinate priming (Dantas et al., 2022; Sogin et al., 2022), although this cannot be established conclusively without direct microbial community profiling. These observations highlight an underappreciated dimension of exogenous succinate application: its potential capacity to reshape the rhizosphere microbiome, which may subsequently influence the process of pea-nodule establishment.

**Figure 7:**
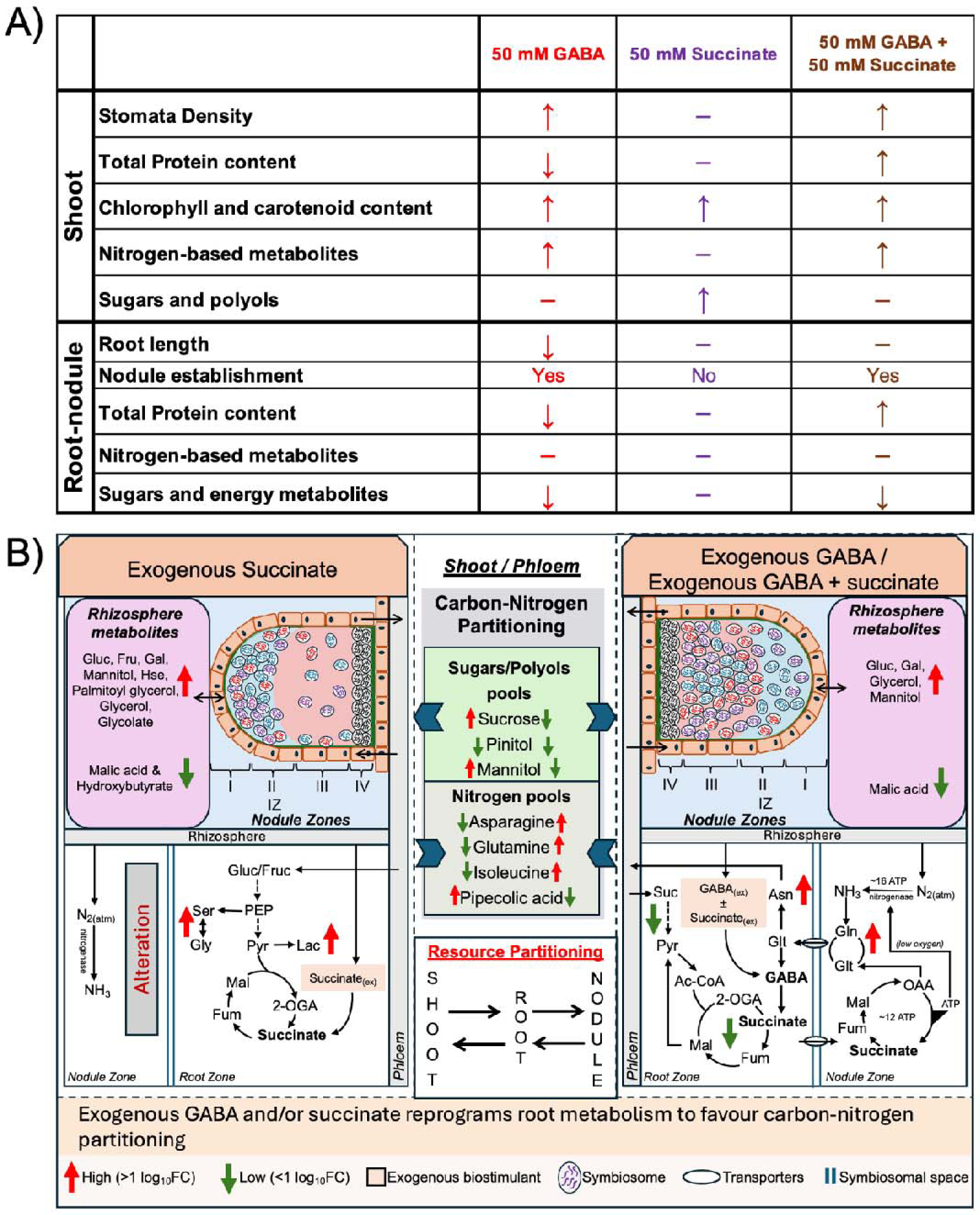
Schematic summary of the observed metabolic phenotypes across treatments. A) Morpho-physiology of the response. Based on the treatments, the response was color-coded. The color codings represent the treatment conditions, and the arrows represent the changes observed. B) Metabolic alterations in the root-nodule compartment. Suc: Sucrose, Gal: Galactose, Gluc: Glucose, Fruc: Fructose, PEP: Phosphoenol-pyruvate, Pyr: Pyruvate, Lac: Lactic acid, Ac-CoA: Acetyl-CoA, 2-OGA: Oxoglutarate, Fum: Fumarate, Mal: Malate, OAA: Oxaloacetate, Glt: Glutamate, Gln: Glutamine, Ser: Serine, Gly: Glycine, Hse: Homoserine. Red upward arrows signify accumulation, and green downward arrows represent consumption of the represented metabolites. FC: Fold change values w.r.t. day 0 control samples.

## 5. Conclusions

The present study demonstrates that rhizospheric application of GABA shunt intermediates induces distinct metabolic reprogramming in pea (*Pisum sativum* var. AS-10) under symbiotic field conditions. Exogenous GABA (50 mM) functioned as a metabolic biostimulant by remodeling carbon-nitrogen partitioning across shoot and nodulated root compartments, evidenced by depletion of TCA cycle intermediates and soluble sugars in nodulated roots alongside substantial accumulation of asparagine (+3.9 log_10_-fold relative abundance) and elevated elemental nitrogen content in shoot tissue (Figure 7). Comparative metabolomics under non-symbiotic conditions confirmed that the nitrogen-assimilatory response to GABA is strictly dependent on an active rhizobium symbiosis, highlighting the context-specificity of GABA biostimulant activity. Multi-platform validation using GC-MS and ^1^H-NMR yielded consistent evidence for the major metabolic shifts reported, corroborating the robustness of the key findings. In contrast, exogenous succinate application disrupted nodule organogenesis and altered rhizosphere metabolite composition without inducing coordinated carbon-nitrogen reprogramming, underscoring the mechanistic specificity of GABA shunt activity relative to direct dicarboxylate supplementation. Collectively, these findings establish exogenous GABA as a promising candidate metabolic biostimulant for modulating carbon-nitrogen dynamics in legume-*rhizobium* symbiosis, with potential implications for sustainable nitrogen management in grain legume production. Future studies incorporating direct nitrogenase activity measurements, ^13^C-GABA metabolic flux analysis, ^15^N_2_ incorporation assays, and dose-response optimisation under field conditions will be necessary to translate these metabolic observations into agronomically applicable biostimulant strategies.

## Contributions

RN-conceptualization, performed experiments, data curation, formal analysis, validation, visualisation, writing-original draft, PDS - methodology, writing-review and editing, SKM-conceptualization, funding acquisition, supervision, review and editing of draft.

## Acknowledgment

RN and PDS acknowledge the Ministry of Education for MTech (by Research) and PhD fellowships, respectively. We acknowledge the FIST confocal microscopy facility at IIT Mandi (File Number: SR/FST/LS-II/2022/950). We would like to acknowledge Dr. Shweta Tandon and Dr. Baskar Bakthavachalu for their support with confocal microscopy.

## Funding

SKM acknowledges ANRF (Anusandhan National Research Foundation) for the support (File no.: ECR/2016/001176)

## Declarations of Interest

### Ethical approval

The study complied with the ethical standards.

### Competing interests

The authors declare no competing interests.

### Data availability

The datasets generated and analysed during the current study are available from the corresponding author on reasonable request.

## Notes

### Competing Interest Statement

The authors have declared no competing interest.

