## Supplementary File for "Decoding central metabolic rewiring induced by exogenous GABA shunt intermediates in Pea"

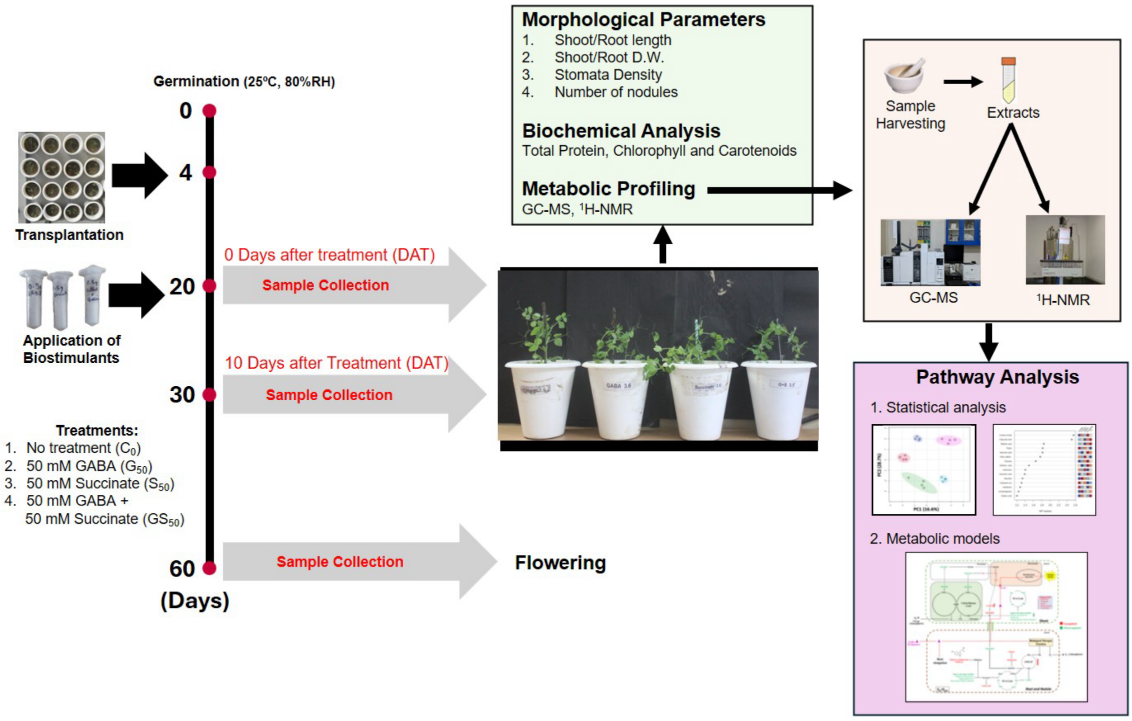


**Supplementary Figure S1: Schematic workflow adopted to define the metabolic phenotypes**.

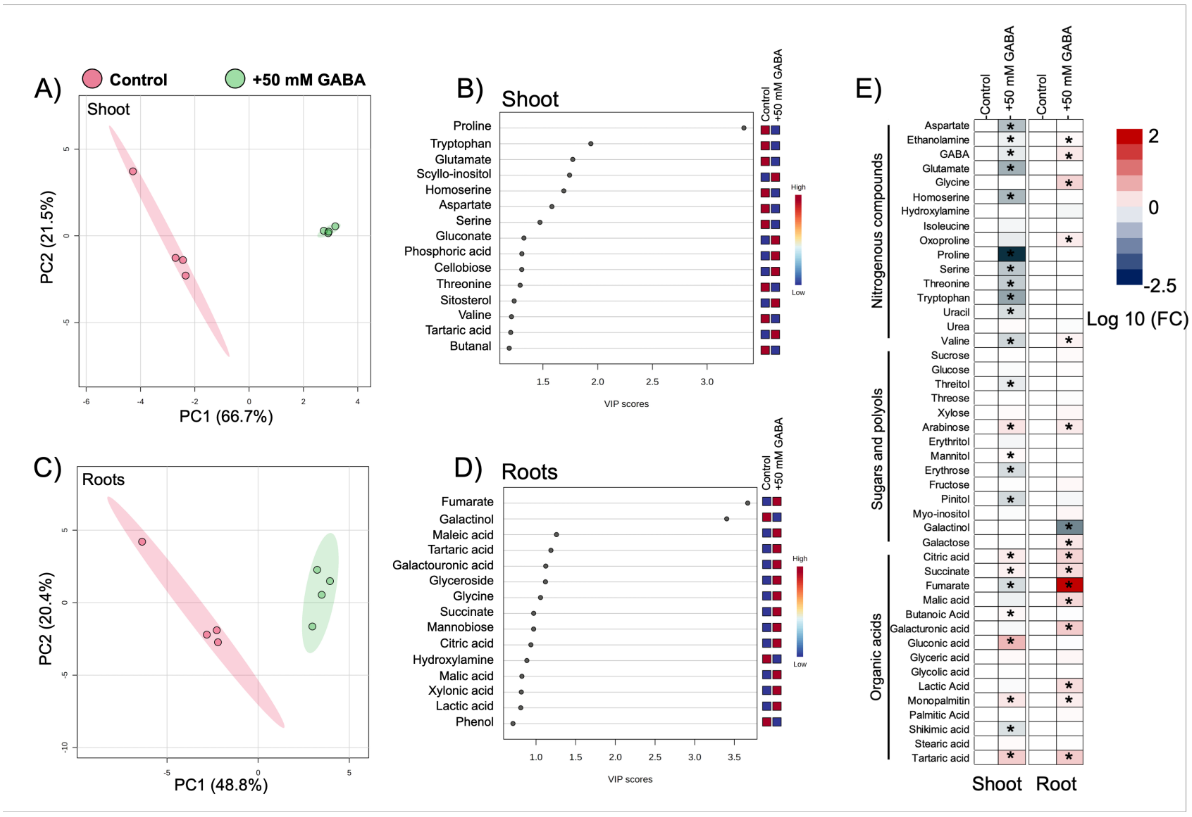


**Supplementary Figure S2: Metabolic changes in the pea plants upon exposure to 50 mM GABA under non-symbiotic conditions**. A-B) Metabolic variations in the shoot upon exogenous treatment, captured by principal component analysis and VIP score plots of the samples. The shaded regions highlight 95% confidence intervals between the treatments. C-D) Metabolic variations in the root under non-symbiotic conditions, upon exogenous treatment, captured by principal component analysis and VIP score plots of the samples. The shaded regions highlight 95% confidence intervals between the treatments. E) Heatmap showcasing the changes in the metabolite levels with respect to control conditions. The values are represented in terms of log 10-fold change compared to the control. The significant values are indicated with an asterisk (p<0.5).

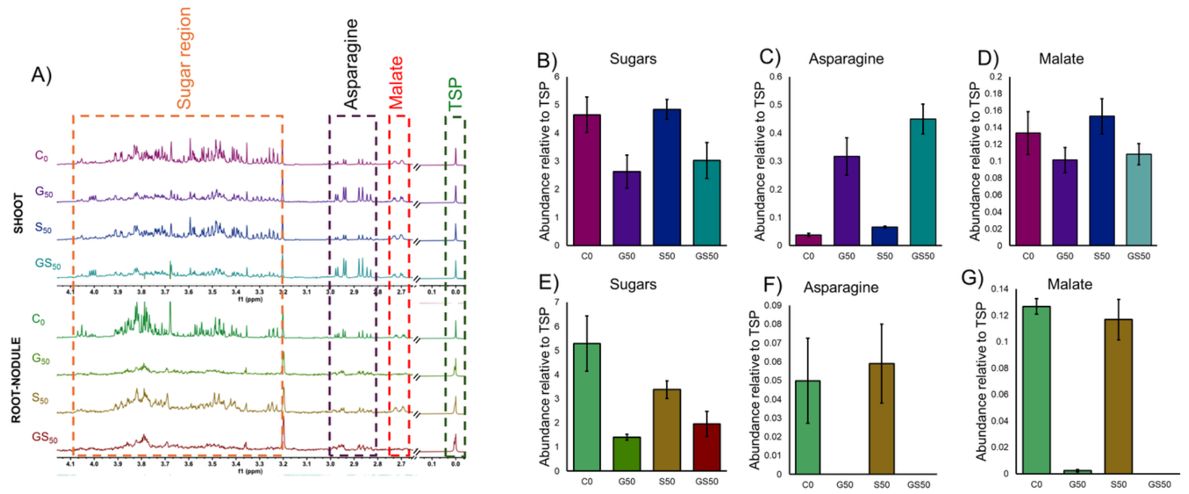


**Supplementary Figure S3: 1H-NMR based metabolic profiling of shoot and nodulated pea root highlights the absolute changes in the concentrations of sugars, asparagine and malate.** A) Spectral representation of the shoot and root-nodule, respectively. The identified metabolites are as follows: Sugar region, Asparagine, malate and 0.01% (w/v) T.S.P as the internal standard. B-D) shows the absolute levels of Sugars, Asparagine and Malate, respectively wrt levels of TSP shows the absolute levels of Sugars, Asparagine and Malate, respectively wrt levels of TSP in the nodulated pea roots. The abbreviations used are control (C0), +50 mM GABA (G50), +50 mM Succinate (S50), +50 mM GABA and +50 mM succinate (GS50), TSP: Trimethylsilyl-2,2,3,3-tetradeuteropropionic acid sodium salt (Internal standard).

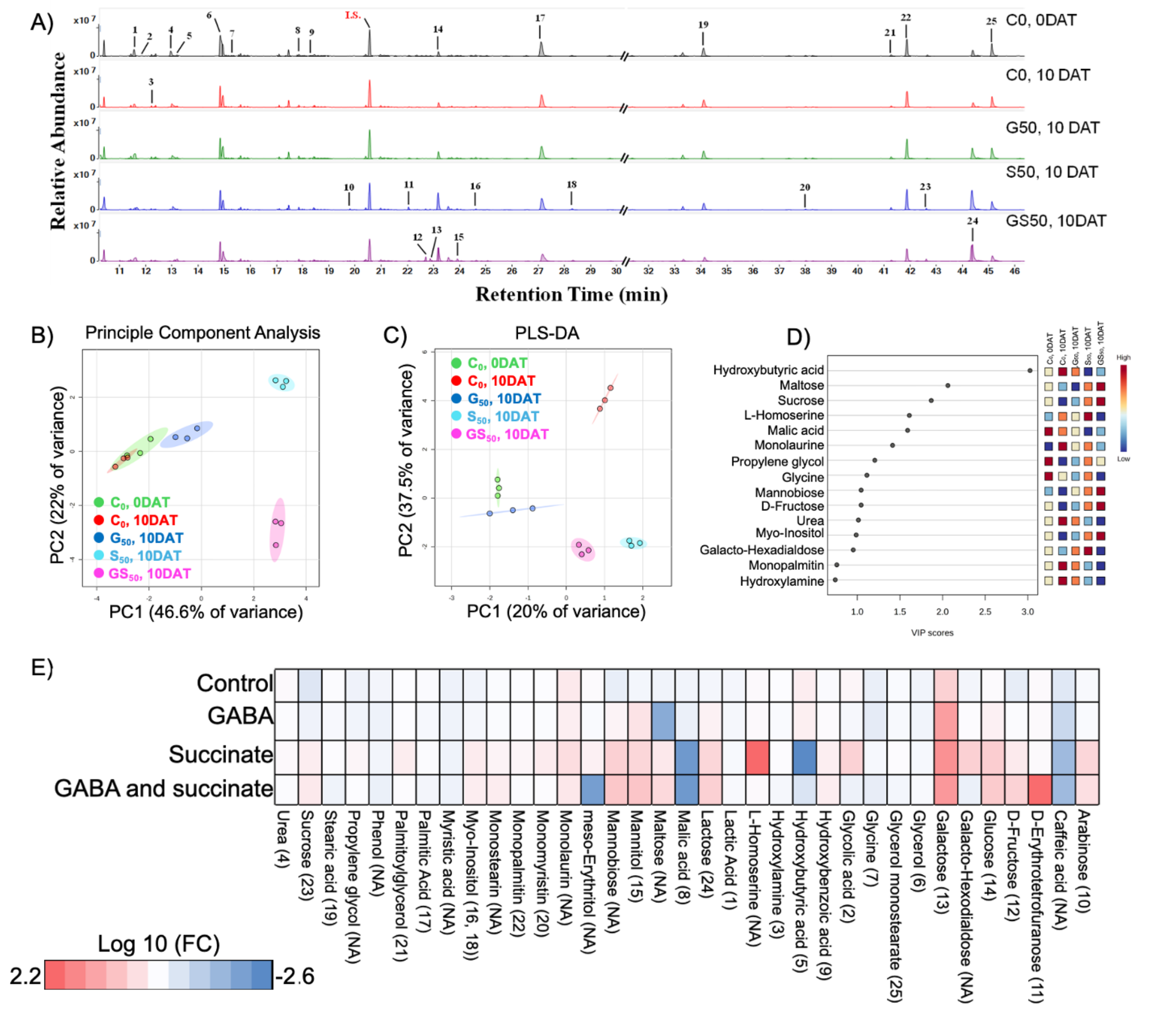


**Supplementary Figure S4: GC-MS based metabolic profiling of soil exudation.** A) GC-MS spectra representing the kinetic variations in soil exudation from 0 DAT to 10 DAT across treatments. The labels annotated represents: 1) lactic acid, 2) glycolic acid, 3) hydroxylamine, 4) Urea, 5) 3-hydroxybutyric acid, 6) glycerol, 7) glycine, 8) malic acid, 9) oxoproline, 10) arabinose, 11) erythrofuranose, 12) fructose, 13) galactose, 14) glucose, 15) mannitol/sorbitol, 16&18) myo-inositol, 17) palmitic acid, 19) stearic acid, 20) monomyristin, 21) palmitoylglycerol, 22) monopalmitin, 23) sucrose, 24) lactose, 25) glycerol monostearate and IS) internal standard. B) represents the variation in the principle component analysis. C-D) PLS-DA and VIP score plots of the samples. E) Heatmap showcasing the relative log 10 fold change values w.r.t. C0, 0DAT. C0, 0 DAT – rhizosphere metabolites without any treatment at day 0; C0, 10DAT – rhizosphere metabolites without any treatment at day 10; G50, 10DAT – rhizosphere metabolites upon treatment with 50 mM GABA at day 10; S50, 10DAT – rhizosphere metabolites upon treatment with 50 mM succinate at day 10; GS50, 10DAT – rhizosphere metabolites upon treatment with 50 mM GABA and 50 mM succinate at day 10.

**Supplementary Table S1. The global metabolite profiles of pea under non-symbiotic conditions: control (C0) and +50 mM GABA (G50).** The values are depicted as mean of sample metabolite profiles, normalised with internal standard.

| Sr. No. | Compound Class | Metabolite name (label on spectra) | C0, Shoot | G50, Shoot | C0, Root | G50, Root |
|---|---|---|---|---|---|---|
| 1 | **TCA Dicarboxylates** | Fumarate **(10)** | 13.9 | 5.2 | 0.0 | 6.7 |
| 2 | | Succinate **(11)** | 60.8 | 77.5 | 8.6 | 15.5 |
| 3 | | Malic acid **(15)** | 239.3 | 161.9 | 83.9 | 157.5 |
| 4 | | Citric acid **(21)** | 147.9 | 204.8 | 120.1 | 241.9 |
| 6 | **Nitrogen Compounds** | Hydroxylamine **(3)** | 0.0 | 0.0 | 21.6 | 16.9 |
| 7 | | Urea **(4)** | 103.1 | 111.2 | 103.7 | 96.2 |
| 8 | | Valine **(5)** | 8.0 | 2.6 | 1.3 | 1.6 |
| 9 | | Ethanolamine **(6)** | 73.9 | 50.5 | 49.7 | 56.3 |
| 10 | | Isoleucine **(7)** | 6.2 | 4.9 | 0.0 | 0.0 |
| 11 | | Proline **(8)** | 33.0 | 0.0 | 0.0 | 0.0 |
| 12 | | Glycine **(9)** | 0.0 | 0.0 | 6.9 | 14.5 |
| 13 | | Uracil **(NA)** | 8.8 | 3.4 | 0.0 | 0.0 |
| 14 | | Serine **(12)** | 100.3 | 21.4 | 0.0 | 0.0 |
| 15 | | Threonine **(13)** | 73.7 | 17.8 | 0.0 | 0.0 |
| 16 | | Homoserine **(14)** | 92.1 | 12.7 | 111.4 | 110.3 |
| 17 | | Aspartic acid **(16)** | 74.4 | 12.6 | 0.0 | 0.0 |
| 18 | | Oxoproline **(17)** | 142.3 | 82.8 | 11.3 | 15.5 |
| 19 | | Aminobutanoic acid **(18)** | 82.1 | 46.4 | 14.4 | 21.3 |
| 20 | | Glutamic acid **(19)** | 122.6 | 13.7 | 0.0 | 0.0 |
| 22 | **Organic Acid and Other compounds** | Lactic Acid **(1)** | 66.1 | 58.8 | 45.9 | 81.7 |
| 23 | | Glycolic acid **(2)** | 7.0 | 7.0 | 0.0 | 0.0 |
| 24 | | Glyceric acid **(NA)** | 72.6 | 80.8 | 19.8 | 23.1 |
| 25 | | Shikimic acid **(20)** | 17.1 | 7.4 | 0.0 | 0.0 |
| 26 | | Gluconic acid **(28)** | 13.3 | 43.7 | 12.1 | 11.8 |
| 27 | | Palmitic Acid **(29)** | 149.1 | 150.7 | 123.7 | 134.5 |
| 28 | | Monopalmitin **(32)** | 44.9 | 66.9 | 24.0 | 31.8 |
| 50 | **Sugar and sugar alcohols** | Erythritol **(22)** | 51.3 | 40.8 | 0.0 | 0.0 |
| 51 | | Fructose **(23)** | 356.2 | 342.0 | 508.1 | 576.5 |

| | | | | | | |
|---|---|---|---|---|---|---|
| **52** | | Glucose **(24)** | 874.4 | 785.7 | 649.0 | 673.6 |
| **53** | | Mannitol/Sorbitol **(25)** | 13.5 | 15.1 | 0.0 | 0.0 |
| **54** | | Pinitol **(26)** | 70.3 | 24.4 | 21.8 | 17.8 |
| **55** | | Mannose **(27)** | 141.8 | 67.2 | 0.0 | 0.0 |
| **56** | | Myo-inositol **(30)** | 381.8 | 355.2 | 182.2 | 214.7 |
| **57** | | Galactose **(31)** | 139.3 | 133.0 | 40.4 | 60.9 |
| **58** | | Sucrose **(33)** | 841.9 | 866.5 | 1154.2 | 1259.9 |

**Supplementary Table S2. The kinetic metabolite profiles of Pea shoot under four conditions: control (C0), +GABA (G50) , +Succinate (S50) and +GABA/+Succinate (GS50).** The values are depicted as mean of sample metabolite profiles, normalised with internal standard.

| Sr. No | Compound Class | Metabolite Name (Label on spectra) | C0, 0DAT | C0, 10DAT | G50, 10DAT | S50, 10DAT | GS50, 10DAT |
|---|---|---|---|---|---|---|---|
| 1 | **TCA Dicarboxylates** | Citric acid **(30)** | 112.8 | 164.8 | 96.8 | 156.0 | 150.1 |
| 2 | | Succinate **(10)** | 37.8 | 32.8 | 17.7 | 26.0 | 23.3 |
| 3 | | Malic acid **(16)** | 196.7 | 290.7 | 236.3 | 225.0 | 227.2 |
| 4 | | Maleic Acid **(9)** | 48.4 | 55.8 | 32.8 | 36.6 | 38.0 |
| 5 | | Fumaric Acid **(13)** | 67.3 | 33.0 | 0.1 | 29.2 | 70.5 |
| 6 | **Nitrogen Based** | Threonine **(8)** | 90.7 | 60.8 | 44.9 | 58.1 | 51.6 |
| 7 | | GABA **(19)** | 84.8 | 73.8 | 54.0 | 57.2 | 58.9 |
| 8 | | Homoserine **(14)** | 417.5 | 307.3 | 273.7 | 264.0 | 262.6 |
| 10 | | Pipecolic acid **(39)** | 208.5 | 78.2 | 15.5 | 173.6 | 31.6 |
| 11 | | Butanoic acid **(NA)** | 12.4 | 24.7 | 10.8 | 22.1 | 10.9 |
| 12 | | Oxoproline (**18)** | 55.2 | 52.2 | 55.0 | 71.6 | 90.6 |
| 13 | | Asparagine **(23)** | 322.5 | 0.1 | 196.5 | 16.3 | 324.6 |
| 14 | | Aspartic acid **(15)** | 115.2 | 105.2 | 95.0 | 70.3 | 101.0 |
| 15 | | Valine **(5)** | 13.0 | 5.8 | 5.6 | 5.1 | 6.5 |
| 16 | | Lysine **(NA)** | 11.9 | 3.5 | 2.6 | 5.2 | 7.4 |
| 17 | | Serine **(6)** | 49.8 | 43.3 | 47.1 | 45.1 | 69.4 |
| 18 | | Isoleucine **(7)** | 7.1 | 6.3 | 8.1 | 0.1 | 8.4 |
| 19 | | Glutamic acid **(25)** | 15.7 | 4.7 | 17.3 | 15.3 | 30.1 |
| 20 | | Urea **(3)** | 84.8 | 71.0 | 79.2 | 86.0 | 95.5 |
| 21 | | Uracil (**12)** | 10.8 | 8.7 | 6.9 | 5.8 | 8.5 |
| 23 | | Ethanolamine **(NA)** | 69.1 | 53.8 | 67.7 | 61.8 | 73.3 |
| 24 | | Hydroxylamine **(2)** | 18.5 | 15.8 | 27.5 | 19.5 | 26.7 |
| 25 | **Organic acids and others** | Palmitic Acid **(37)** | 118.9 | 68.2 | 92.5 | 95.3 | 109.0 |
| 26 | | Glyceric acid **(11)** | 88.0 | 161.9 | 71.7 | 102.7 | 68.7 |
| 27 | | Glycerol monostearate **(NA)** | 18.1 | 5.3 | 6.0 | 23.9 | 16.9 |
| 28 | | Glycolic acid **(1)** | 7.4 | 7.6 | 5.6 | 5.8 | 5.6 |

| | | | | | | | |
|---|---|---|---|---|---|---|---|
| **29** | | Hexanoic acid **(NA)** | 17.6 | 12.2 | 8.2 | 23.3 | 10.0 |
| **30** | | Ribonic acid **(NA)** | 98.1 | 82.7 | 89.5 | 72.8 | 101.9 |
| **31** | | Ritalinic acid **(NA)** | 326.7 | 92.3 | 49.7 | 104.4 | 82.3 |
| **32** | | Shikimic acid **(29)** | 7.7 | 0.1 | 4.2 | 8.3 | 6.6 |
| **33** | | Stearic acid **(NA)** | 43.3 | 17.1 | 30.6 | 33.6 | 39.1 |
| **34** | | Galacturonic acid **(NA)** | 10.3 | 12.7 | 8.9 | 19.8 | 12.6 |
| **35** | | Gluconic acid **(NA)** | 22.9 | 3.9 | 15.6 | 79.4 | 27.6 |
| **36** | | Lactic Acid **(NA)** | 33.7 | 36.6 | 45.7 | 42.7 | 53.1 |
| **37** | | Monopalmitin **(NA)** | 28.8 | 12.8 | 22.4 | 29.1 | 29.8 |
| **38** | | Phenol **(NA)** | 1.3 | 0.6 | 0.8 | 1.1 | 0.9 |
| **39** | | Galactaric acid **(NA)** | 2.7 | 2.7 | 2.3 | 2.0 | 3.7 |
| **40** | | Phosphoric acid **(4)** | 10.7 | 3.2 | 1.9 | 4.9 | 7.7 |
| **41** | | 3-Amino-2-piperidone **(17)** | 186.2 | 0.1 | 60.7 | 0.1 | 208.2 |
| **42** | **Sugars and sugar alcohols** | Fructose **(32)** | 198.7 | 370.1 | 319.7 | 549.5 | 313.9 |
| **43** | | Threose **(21)** | 41.7 | 37.9 | 29.0 | 43.0 | 25.7 |
| **44** | | Xylose **(20)** | 161.4 | 179.0 | 200.6 | 260.0 | 182.2 |
| **45** | | Sucrose **(30)** | 325.0 | 434.9 | 435.9 | 424.9 | 375.8 |
| **46** | | Glucose **(34)** | 1006.2 | 893.1 | 684.7 | 985.8 | 371.3 |
| **47** | | Arabinofuranose **(28)** | 157.1 | 134.6 | 165.3 | 206.0 | 134.2 |
| **48** | | Arabinose **(26)** | 162.9 | 37.9 | 17.0 | 32.4 | 95.7 |
| **49** | | Melibiose **(41)** | 13.9 | 7.1 | 5.7 | 37.9 | 14.9 |
| **50** | | Mannobiose **(NA)** | 30.4 | 28.2 | 8.6 | 64.7 | 28.6 |
| **51** | | Erythro-Pentitol **(22)** | 5.7 | 5.3 | 3.6 | 6.2 | 0.1 |
| **52** | | Mannitol **(35)** | 11.3 | 61.3 | 8.2 | 5.2 | 0.1 |
| **53** | | Galactinol **(NA)** | 0.1 | 2.9 | 0.1 | 7.7 | 0.1 |
| **54** | | 2-O-Glycerol-ɑ-d-galactopyranoside **(NA)** | 75.8 | 59.7 | 64.4 | 106.7 | 77.9 |
| **55** | | Anhydroglucitol **(31)** | 39.3 | 41.8 | 24.9 | 38.7 | 18.2 |
| **56** | | Levoglucosan **(NA)** | 40.1 | 41.7 | 34.3 | 42.8 | 23.0 |
| **57** | | Methyl xylopyranoside **(NA)** | 6.7 | 4.4 | 3.6 | 7.9 | 5.4 |
| **58** | | Myo-inositol **(38)** | 252.3 | 447.1 | 236.8 | 461.1 | 252.9 |
| **59** | | Cellobiose **(NA)** | 27.4 | 18.8 | 15.9 | 8.2 | 41.4 |

| | | | | | | | |
|---|---|---|---|---|---|---|---|
| **60** | | Galactose **(33)** | 277.2 | 267.5 | 76.0 | 178.4 | 457.8 |
| **61** | | Ethyl-α-D-glucopyranoside **(NA)** | 54.8 | 48.2 | 120.0 | 105.9 | 150.8 |
| **62** | | Pinitol **(36)** | 19.4 | 32.8 | 27.4 | 36.7 | 45.9 |

**Supplementary Table S3. The kinetic metabolite profiles of Pea root-nodule under four conditions: control (C0), +GABA (G50), +Succinate (S50) and +GABA/+Succinate (GS50).** The values are depicted as the mean of sample metabolite profiles, normalised with the internal standard.

| Sr. No. | Compound Class | Metabolite name (Label on spectra) | C0, 0DAT | C0, 10DAT | G50, 10DAT | S50, 10DAT | GS50, 10DAT |
|---|---|---|---|---|---|---|---|
| 1 | **TCA Dicarboxylates** | Citric acid **(24)** | 58.1 | 164.5 | 37.4 | 92.7 | 53.1 |
| 2 | | Maleic acid **(11)** | 0.1 | 39.5 | 0.1 | 0.1 | 0.1 |
| 3 | | Fumarate **(9)** | 8.8 | 23.8 | 6.8 | 11.2 | 5.3 |
| 4 | | Malic acid **(14)** | 164.8 | 336.8 | 158.6 | 270.6 | 159.3 |
| 5 | | Succinate **(10)** | 17.0 | 18.0 | 11.8 | 14.3 | 12.8 |
| 6 | **Nitrogen Compounds** | Asparagine **(18)** | 48.2 | 16.4 | 7.1 | 12.4 | 0.1 |
| 7 | | Aspartate **(NA)** | 14.0 | 0.1 | 0.1 | 0.1 | 0.1 |
| 8 | | Ethanolamine **(6)** | 40.6 | 65.9 | 47.3 | 57.8 | 53.4 |
| 9 | | GABA **(16)** | 28.2 | 48.0 | 15.2 | 23.7 | 23.3 |
| 10 | | Glycine **(8)** | 22.2 | 0.1 | 20.5 | 28.1 | 18.5 |
| 11 | | Homoserine **(12)** | 885.6 | 330.5 | 386.3 | 631.5 | 488.2 |
| 12 | | Hydroxylamine **(2)** | 17.9 | 17.8 | 20.5 | 20.4 | 19.9 |
| 13 | | Oxoproline **(15)** | 21.8 | 0.1 | 0.1 | 10.6 | 0.1 |
| 14 | | Putrescine **(22)** | 15.8 | 5.5 | 5.6 | 3.6 | 3.4 |
| 15 | | Serine **(5)** | 20.5 | 7.8 | 0.1 | 12.1 | 0.1 |
| 16 | | Threonine **(7)** | 76.3 | 25.0 | 28.9 | 35.6 | 35.6 |
| 17 | | Uracil **(NA)** | 11.1 | 6.3 | 0.1 | 6.4 | 0.1 |
| 18 | | Urea **(3)** | 93.8 | 117.9 | 70.2 | 92.4 | 74.4 |
| 19 | | Valine **(4)** | 5.4 | 0.1 | 0.1 | 3.0 | 0.1 |
| 20 | **Organic acids and other Compounds** | Butanoic acid **(13)** | 11.4 | 21.1 | 5.4 | 37.3 | 7.1 |
| 21 | | Galactaric acid **(NA)** | 2.2 | 3.6 | 0.1 | 0.1 | 0.1 |

| | | | | | | | |
|---|---|---|---|---|---|---|---|
| **22** | | Galacturonic acid **(36)** | 9.2 | 14.5 | 5.9 | 12.2 | 6.0 |
| **23** | | Gluconic Acid **(32)** | 46.1 | 95.5 | 43.8 | 70.7 | 15.5 |
| **24** | | Glycerol monostearate **(41)** | 11.0 | 14.5 | 6.9 | 8.6 | 3.8 |
| **26** | | Lactic Acid **(1)** | 31.7 | 45.4 | 72.6 | 58.5 | 62.6 |
| **27** | | Monopalmitin **(39)** | 25.4 | 30.0 | 24.8 | 24.5 | 30.3 |
| **28** | | Palmitic Acid **(33)** | 136.9 | 145.9 | 127.9 | 139.6 | 140.2 |
| **29** | | Phosphoric acid | 1.5 | 0.1 | 0.1 | 1.4 | 0.1 |
| **30** | | Ribonic acid **(23)** | 37.1 | 42.5 | 13.4 | 22.3 | 10.9 |
| **31** | | Stearic acid **(35)** | 54.1 | 55.0 | 51.6 | 54.8 | 67.0 |
| **32** | | Tartaric acid **(19)** | 80.5 | 98.6 | 42.9 | 116.5 | 55.7 |
| **33** | | Trihydroxybutyric acid **(17)** | 20.5 | 19.7 | 5.2 | 24.0 | 7.4 |
| **35** | | Mannobiose **(37)** | 80.7 | 147.0 | 39.2 | 129.5 | 43.4 |
| **36** | | Anhydroglucitol **(NA)** | 9.5 | 11.1 | 5.0 | 10.0 | 4.1 |
| **37** | | Cellobiose **(38)** | 6.7 | 82.9 | 39.0 | 7.9 | 2.4 |
| **38** | | Ethyl-α-D-glucopyranoside **(NA)** | 15.5 | 18.7 | 4.4 | 11.4 | 0.1 |
| **39** | | Fructose **(25)** | 340.6 | 699.9 | 280.1 | 604.7 | 395.2 |
| **40** | | Galactose **(26)** | 28.1 | 57.7 | 4.1 | 52.8 | 10.9 |
| **41** | | Glucose **(27)** | 365.7 | 435.0 | 238.4 | 498.6 | 302.4 |
| **42** | | Levoglucosan **(21)** | 9.6 | 10.5 | 4.2 | 8.3 | 0.1 |
| **43** | | Mannitol **(29)** | 10.6 | 72.1 | 11.1 | 19.8 | 4.2 |
| **44** | | Mannose **(28)** | 145.3 | 273.0 | 43.9 | 308.9 | 77.2 |
| **45** | | Pinitol **(30)** | 52.9 | 181.0 | 52.8 | 80.6 | 41.2 |
| **46** | | Sucrose **(40)** | 444.7 | 467.4 | 324.3 | 459.8 | 253.4 |

| | | | | | | | |
|---|---|---|---|---|---|---|---|
| **48** | | Myo-Inositol **(34)** | 222.4 | 269.2 | 124.2 | 244.8 | 84.1 |
